# Quantifying Cancer Drug Targetability across Biochemical Pathways

**DOI:** 10.1101/2024.09.27.615497

**Authors:** Stuart Hagler

**Affiliations:** Division of Bioinformatics and Computational Biology, Department of Medical Informatics and Clinical Epidemiology, Oregon Health & Science University, Portland, OR 97239, USA; Oregon Clinical and Translational Research Institute, Oregon Health & Science University, Portland, OR 97239, USA

## Abstract

The evaluation of cancer drug treatments within the complex context of biological networks and pathways is challenging. Drugs can be “promiscuous,” targeting multiple genes or pathways, not just the ones of interest. This complexity makes it difficult to assess the full impact of a drug solely based on individual knowledgebases. We make the impact more comprehensible through the characterization of drug targetability in terms of a combination of biochemical pathway data from the Reactome Knowledgebase and drug-target interaction data from the Cancer Targetome. This leads to a graphical representation of drug targetability. We further improve comprehensibility by developing metrics to quantify the graph structures in order to evaluate drug target specificity and navigate the hierarchical nature of biological networks and identify candidate treatment options to increase specificity and minimize off-target effects.. We have developed software that calculates these metrics for drugs in the Cancer Targetome. The software is available for research purposes only at https://github.com/sehagler/drug_targetability.

## 1 Introduction

Cancer was the second leading cause of death in the US in 2019 with 599,601 recorded deaths or a rate of 146.0 per 100,000 people.[1] Recent approaches to the treatment of cancer have pivoted from ’one size fits all’ to individualized approaches collected under the banner of precision medicine in which multiple data sources ranging from genomic sequencing to digital health metrics are harnessed to make clinical and therapeutic decisions.[2] A key challenge is assessment of evidence to support therapeutic selection based on molecular characterization of the tumor (mutation, gene expression etc). Target-Drug evidence can aid in this decision making especially in the context of the underlying pathways. Two key resources for this are the Reactome Knowledgeable and the Cancer Targetome.

The Reactome Knowledgebase provides manually curated molecular details of physiological and pathological processes in humans.[3] It is organized as a graph where the vertices are (biochemical) pathways each consisting of a series of connected (biochemical) reactions and the edges denote a parent-child relationship between two pathways indicating that the child pathway contains a subset of the reactions in the parent pathway. Conventionally the aggregate of all these processes in the human body is called the *Reactome*. The graphical structure defined by the Reactome Knowledgebase enables oncologists to not merely understand the protein’s action in individual processes within the graph, but also to quantify the protein’s aggregate action across the entire Reactome. We can think of the pathways as the sets of proteins involved in the corresponding reactions and the edges indicating superset-subset relationships among those sets.

The Cancer Targetome provides a manually curated database of drug-target interactions.[4] For our immediate purpose, we can think of this database as linking drugs to the set of proteins they target. We can think of the aggregate effect of a drug across the Reactome in terms of the combined aggregate actions of the proteins targeted by the drug.

In precision oncology, these resources allow an oncologist to assess: (1) whether a key gene mutated in a patient is targetable (are there drug-gene target relationships in targetome) and (2) whether, for the pathway(s) this gene is a member of, there are other genes that are targetable. The aggregate effect of each drug across the Reactome may be calculated by looking at the combined aggregate actions of the proteins targeted by that drug—both those genes of interest (mutated, overexpressed etc) and the others that share pathways with those genes. This collection of pathways determines a subgraph within the Reactome graph. At this point the oncologist encounters a problem. The subgraph characterizing the aggregate effect of the proteins across the Reactome will be large and complicated. The oncologist requires a method to visualize and summarize across this graph to assess targetabilithy in the context of the specific patient.

In this paper, we aim to provide mathematical basis for reasoning about the aggregate effects of a drug on the Reactome. We do this by characterizing a drug in terms of the proteins targeted by the drug and constructing a subgraph of the reactome consisting of only those pathways containing proteins targeted by the drug. We then construct a set of metrics that can be used to provide information about the size and topology of the subgraphs that characterize the drug.

## 2 Preliminaries

We begin by defining terminiology that we use in the analysis of the Reactome graph as well as auxiliary vertices that we add to the graph and auxiliary sets that we use in constructiong the analysis.

### 2.1 Terminology

#### Augment Vertex

Given a graph Γ, an augmented graph is an auxiliary graph to Γ constructed by making a copy of Γ and adding one or more augmented vertices *V* ^′^ and necessary edges such that each *V* ^′^ is a child vertex to some vertex *V* ∈ Γ (not necessarily the same *V*) and each *V* ^′^ has no child vertices of its own.

#### Augmented Tree

Given a tree *T*, an augmented tree is an auxiliary tree constructed from *T* through the addition of one or more augment vertices.

#### Child Vertex

Given a graph Γ vertex *V* ∈ Γ, a vertex *V* ^′^ ∈ Γ is a child of *V* if the two vertices are connected by an edge and the edge indicates that *V* is the parent and *V* ^′^ is the child.

#### Conjoined Paths

Given a graph Γ, two paths Π, Π^′^ ⊆ Γ are conjoined if they they havve at least one non-root non -⊥ vertex in common.

#### Conjointness

A broad property of a graph Γ reflecting the extent to which the paths in Γ are conjoined. This property may be quantified in various ways. We propose one way of quantifyiing this property in this paper.

#### Crosstalk Lattice

A lattice Λ is a crosstalk lattice if every path Π ⊆ Λ is a crosstalk path.

#### Crosstalk Vertex

Given a graph Γ, a vertex *V* ∈ Γ is a vertex having two or more parent vertices.

#### Edge

A line in a graph connecting two vertices which designates one of the vertices as the parent and the other vertex as the child.

#### Graph

A set of vertices {*V*} and edges *E* such that every edge in *E* connects two different vertices in {*V*} and every vertex in {*V*} is connected to another vertex in {*V*} by an edge in *E*. Furthermore, a graph has no cycles in the sense that we cannot construct a sequence of vertices in {*V*} connected in sequence by edges in *E* where, moving always from parent to child, we return to the original vertex.

#### Lattice

A graph having exactly one root vertex and one leaf vertex.

#### Leaf Vertex

A vertex having no child vertices

#### Lineage Vertex

A vertex whose parent is a root vertex. We denote the set of lineages vertices in a tree by {*L*}, and the number of lineage vertices in a tree by |*L*|.

#### Parent Vertex

Given a graph Γ vertex *V* ∈ Γ, a vertex *V* ^′^ ∈ Γ is a parent of *V* if the two vertices are connected by an edge and the edge indicates that *V* is the child and *V* ^′^ is the parent.

#### Path

Given a graph Γ a path Π ⊆ Γ is a sequence of vertices *V* ∈ Γ, connected by edges *E* ∈ Γ, such that (1) every consecutive pair of vertices in the sequence is connected by an edge designating the first vertex as the parent and the second as the child, (2) the first vertex is a root vertex of Γ, and (3) the last vertex is a leaf vertex of Γ. By our definitions a path is a graph, a lattice, and a tree. We denote the set of paths in a tree by {Π}, and the number of paths in a tree by |Π|.

#### Path Depth

The number of edges in a path.

#### Root Vertex

A vertex having no parent vertices.

#### Subgraph

Given a graph Γ, a subgraph Γ^′^ is a graph whose set of vertices is a subset of the set of vertices in Γ and whose edges are a subset of the edges in graph Γ. This is denoted by Γ^′^ ⊆ Γ.

#### Subtree

A subgraph that is also a tree.

#### Tree

A graph having exactly one root vertex and no crosstalk vertices.

#### Vertex

A point on a graph and conntected to another vertex via an edge. We denote the set of vertices in a tree by {*V* }, and the number of verticess in a tree by |*V* |.

#### 2.1.1 Auxiliary Vertices

#### BOTTOM

A vertex added to the Reactome graph that is the child of all leaf vertices in the original graph, also denoted by ⊥.

#### TOP

A vertex added to the Reactome graph that is the parent of all root vertices in the original graph, also denoted by ⊤

#### 2.1.2 Auxiliary Sets

*𝔇 𝔯 𝔘 𝔤 𝔰* = The set of all drugs in the Cancer Targetome

*𝔅 𝔞 𝔱 𝔥 𝔴 𝔞 𝔶 𝔑 𝔞 𝔴* 𝔢 𝔰 = The set of all pathway names in Reactome Knowledgebase together with ‘Universal Pathway’ and ‘Empty Pathway’

*𝔅* 𝔯 𝔬 𝔱 𝔢 𝔦 𝔫 𝔰 = The set of all proteins in the Reactome Knowledgebase

*𝔖 𝔱 𝔞 𝔟 𝔩 𝔢 ℑ 𝔇 𝔰* = The set of all stable identifiers in Reactome Knowledgebase together with ‘TOP’ and ‘BOTTOM’

## 3 Analysis of the Reactome graph

The Reactome graph provides a description of the biochemical processes in a human being in the form of a structure containing (1) vertices corresponding to (biochemical) pathways each of which represents a series of connected (biochemical) reactions, (2) edges indicating parent-child relationships between vertices where a parent vertex includes all of the reactions contained in any of its child vertices. The strategy we adopt is to characterize the targetability of a drug by identifying subgraphs of the Reactome graph that contain those pathways that include reactions which involve proteins targeted by the drug. In this section, we develop an analysis of the Reactome graph into substructures of that graph that will facilitate this graphical understanding of drug targetability.

### 3.1 Reactome Lattice & Reactome Data Function

We are given Reactome Knowledgebase in the form of a graph. We find that the development of our analysis is smoother if we work with the closely related—auxiliary—Reactome Lattice ℜ_*L*_. We define a *lattice* to be a graph having exactly one root vertex and one leaf vertex. We construct ℜ_*L*_ by making a copy of the Reactome graph and (1) adding a vertex ⊤ (*top*) together with necessary edges so that ⊤ is the parent of all root vertices in the Reactome graph, (2) adding a vertex ⊥ together with necessary edges so that ⊥ (*bottom*) is the child of all leaf vertices in the Reactome grapg. The vertex ⊤ is the *Universal Pathway* containing all reactions, while the vertex ⊥ is the *Empty Pathway* containing no reactions. The point in adding ⊤ is to ensure that the graph we are talking about does not consist of multiple unconnected graphs, and to ensure that every graph we work with has a single root vertex. The ⊥ vertex is added to make the whole graph a lattice. We illustrate what *R*_*L*_ looks like using a much simpler toy model in Fig. 1.

**Figure 1.**
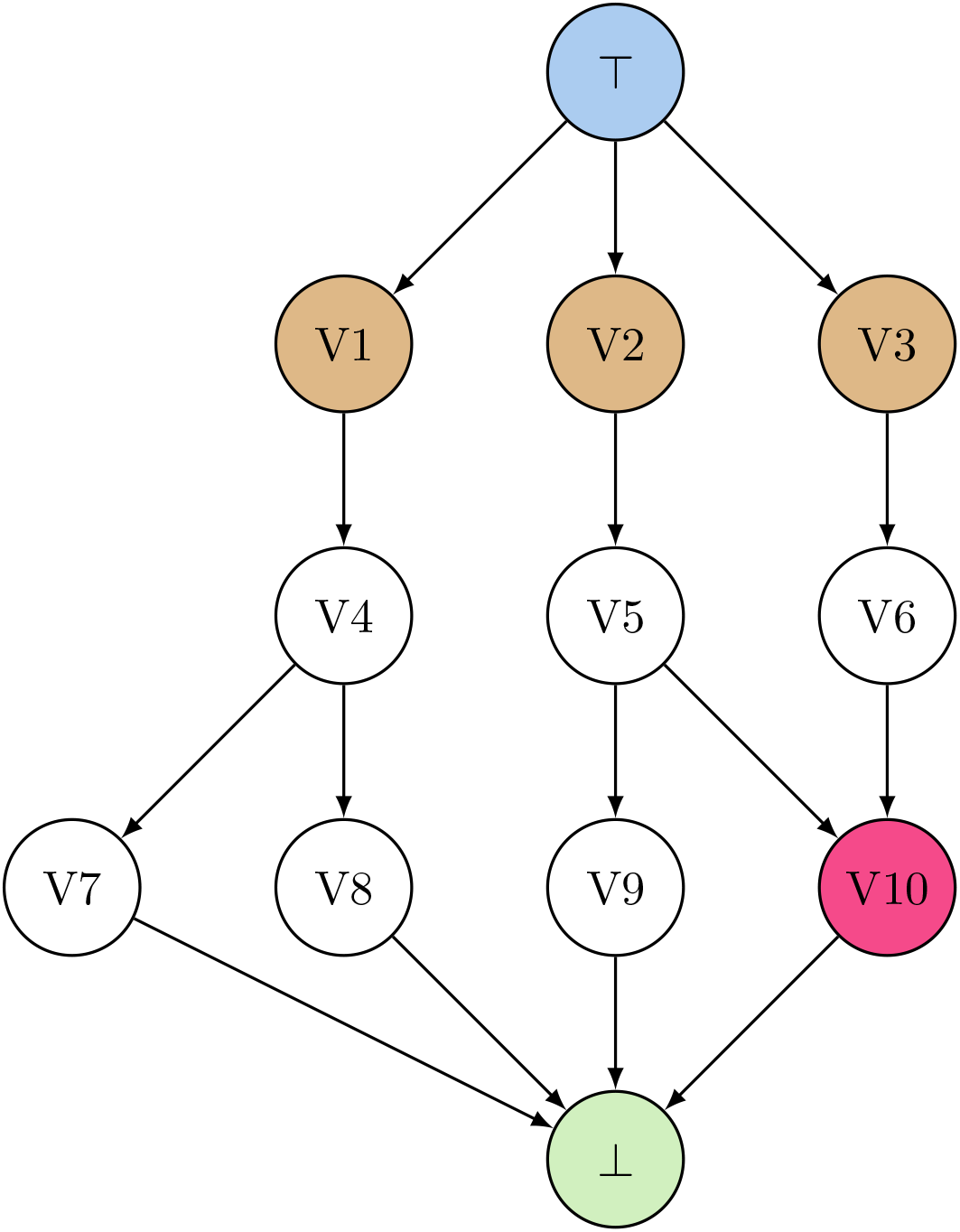
Toy Model Lattice A simple toy lattice. The blue vertex is the root vertex, the brown vertices are lineage vertices, the green vertex is the leaf vertex, the red vertex is a crosstalk vetex, the remaining vertices are white. The edges are indicated by the arrows pointing from parent vertex to child vertex.

The vertices of ℜ_*L*_ are associated with sets of data derving from their association with pathways defined in the Reactome Knowledgebase. We capture this formally by defining function *Reactome*(·) which maps vertices of ℜ_*L*_ to a set of data associated with the vertex. We require that the data set contain the name of the pathway corresponding to the vertex:

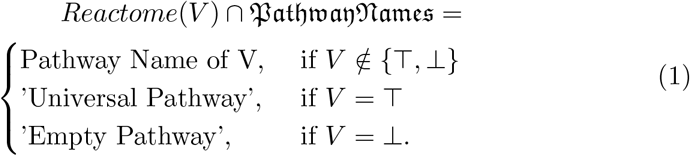

We further require that the data set contain the Reactome stable ID associated with the vertex:

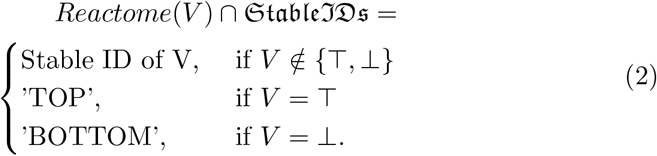

A pathway in the Reactome Knowledgebase is a grouping of interlinked reactions where a reaction is a process that convertes input molecules and/or complexes into output molecules and/or complexes.[5] This means that each pathway is associated with the set of proteins involved in the constituent reactions. We require as well that the data set contain the associated set of proteins:

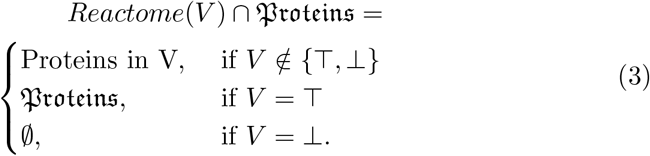

Finally, a pathway contains all of the reactions constituting its child pathways. Thus for two vertices *V, V* ^′^ where *V* ^′^ is a child of *V*, we find:

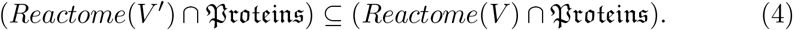

### 3.2 Paths

Our analysis of the Reactome graph is built around a particular graph structure called a *path*. A path is any sequence of vertices beginning with a root vertex and ending with a leaf vertex where each vertex (excepting the first, root vertex in the sequence) is a child of the preceding vertex together with the corresponding edges indicating the parent-child relationship. We illustrate the set of all paths in ℜ_*L*_ using the toy model in Fig. 2.

**Figure 2.**
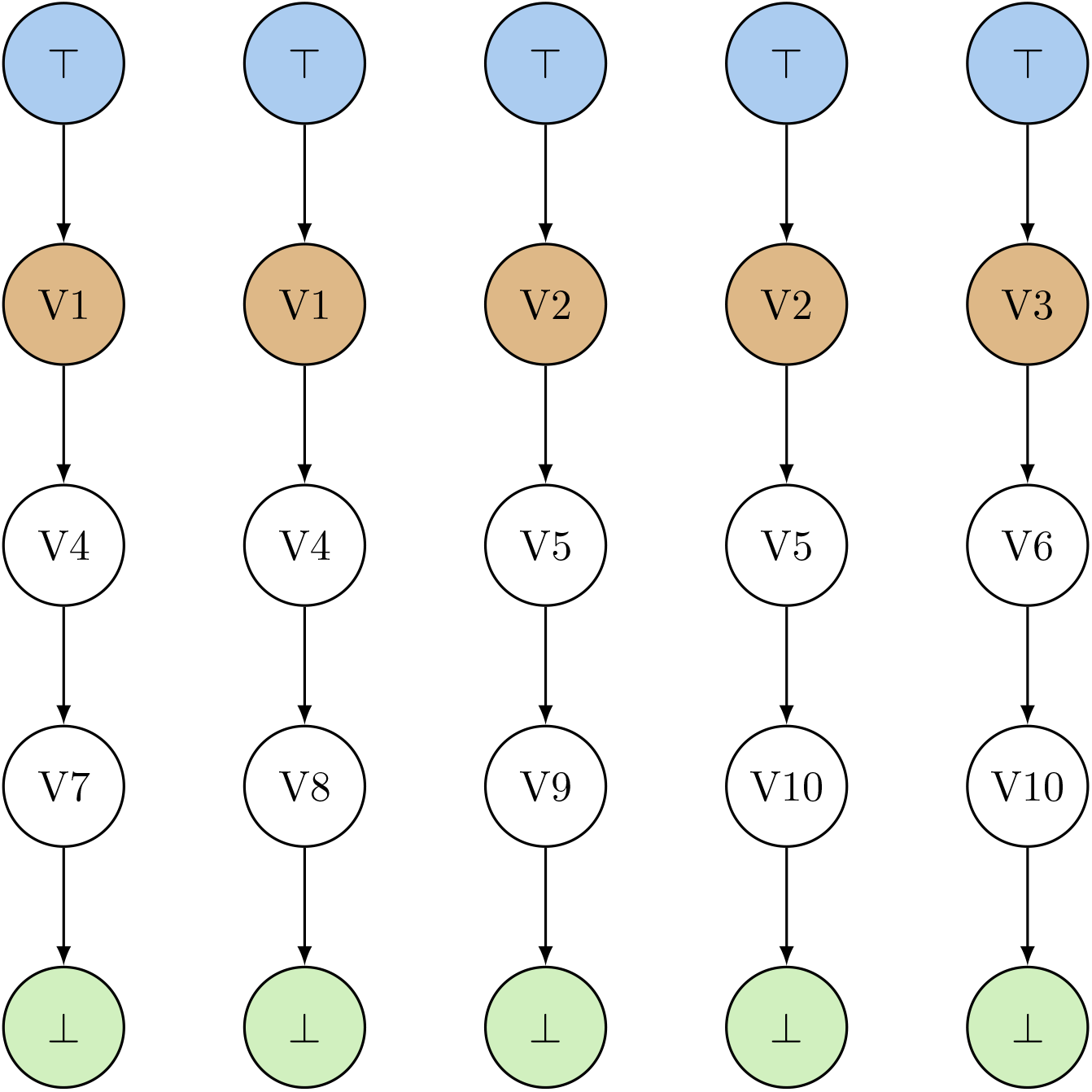
Toy Model Lattice Paths The paths for the toy lattice in Fig. 1. The blue vertices are the root vertices, the brown vertices are lineage vertices, the green vertices are the leaf vertices, the remaining vertices are white. The vertex V10 is not shown as a crosstalik vertex since in these graphs it has only one parent in each graph. Otherwise the vertices are colored the same as in Fig. 1.

We can also think of a path in terms of the sequence of pathways corresponding to the vertices in the path. An example path with root, lineage and leaf vertices explicitly indicated is:

TOP Universal Pathway **(Root)**

R-HSA-112316 Neuronal System **(Lineage)**

R-HSA-112315 Transmission across Chemical Synapses

R-HSA-112314 Neurotransmitter receptors and postsynaptic signal transmission

R-HSA-181431 Acetylcholine binding and downstream events

R-HSA-622327 Postsynaptic nicotinic acetylcholine receptors

R-HSA-629587 Highly sodium permeable postsynaptic acetylcholine nicotinic receptors

BOTTOM Empty Pathway **(Leaf)**

Pathways are groupings of interlinked reactions. At the reaction level, we can interpret a path as a process of zooming into smaller and smaller sets of reactions. As we are more specifically interested in the proteins associated with the reactions, the path is a zooming through smaller and smaller sets of proteins. That is, for a path Π ⊆ ℜ_*L*_ with a vertex sequence *V*_0_, *V*_1_, …, *V*_*N*_, we observe:

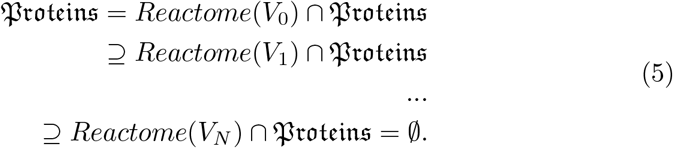

### 3.3 Decomposition of the Reactome Lattice

We can think of the Reactome lattice *R*_*L*_ as a set of paths together with a procedure for determining which vertices are shared by multiple paths in the lattice structure. We define a *crosstalk vertex* to be any non-⊥ vertex having more than one parent. We can then decompose the lattice *R*_*L*_ into two lattices. We do this by first decomposing the set of paths making up *R*_*L*_ into two sets —the set of all paths not containing crosstalk vertices and the set of paths containing crosstalk vertices. We can then construct two lattices from the two sets of paths using the procedure of identifying vertices within the lattice that we have assumed given. We call the lattice constructed from paths not containing crosstalk vertices the *trees lattice*, and the lattice constructed from paths containing crosstalk vertices the *crosstalk lattice*. We define a *tree* to be a graph having exactly one root vertex. We find that if we take the trees lattice and remove the ⊤ and ⊥ vertices together with any edges involving those vertices that we end up with a set of trees. At the end of this procedure we are left with a set of trees and a crosstalk lattice. We illustrate the end result of this process of decomposition for the toy model in Fig. 3.

**Figure 3.**
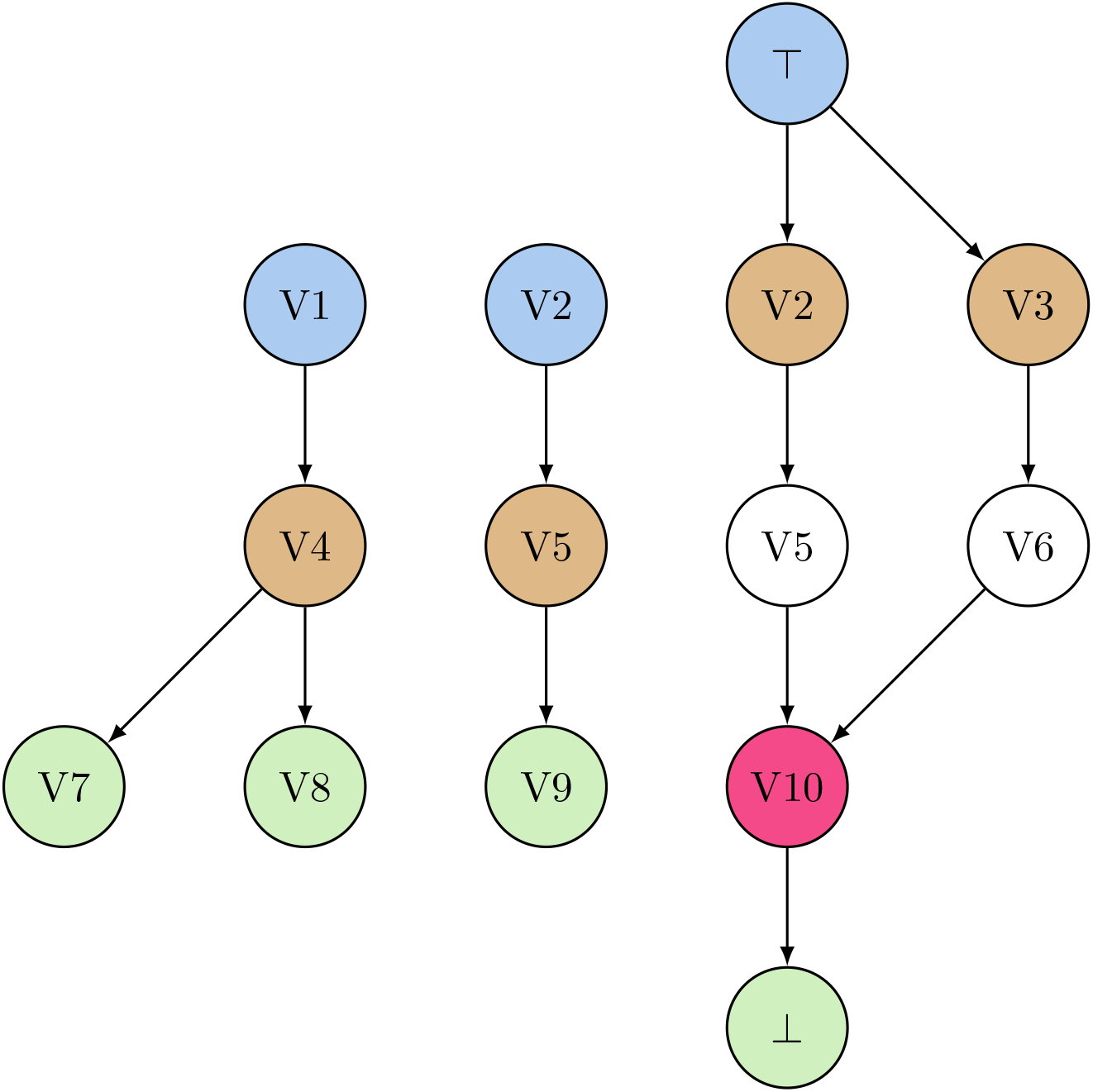
Toy Model Trees & Crosstalk Lattice The trees and crosstalk lattice for the toy model in Fig. 1. The left and center graphs are trees while the right graph is the crosstalk lattice. The blue vertices are the root vertices, the brown vertices are lineage vertices, the green vertices are the leaf vertices, the red vertex is a crosstalk vetex, the remaining vertices are white. The colors of the vertices differ from 1 according to their role in the graphs shown here.

The trees and crosstalk lattice may be further decomposed into paths as above. We illustrate this decomposition into paths in Fig. 4 We give two example paths from the trees we have derived. The first is a path from the R-HSA-162582 tree:

**Figure 4.**
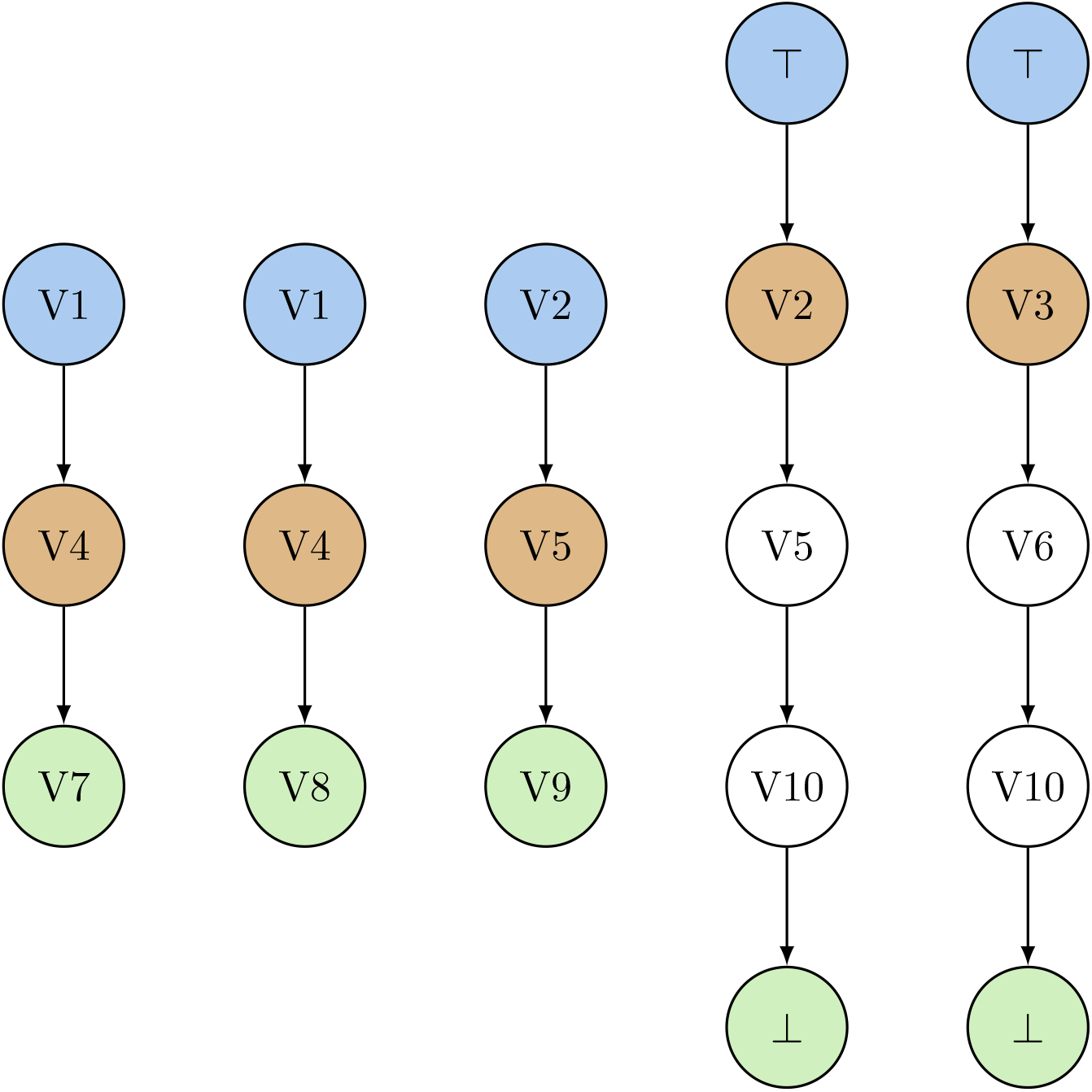
Toy Model Tree & Crosstalk Lattice Paths The paths for the trees and crosstalk lattice in Fig. 3. The blue vertices are the root vertices, the brown vertices are lineage vertices, the green vertices are the leaf vertice, the remaining vertices are white. The vertex V10 is not shown as a crosstalik vertex since in these graphs it has only one parent in each graph. Otherwise the vertices are colored the same as in Fig. 3.

R-HSA-162582 Signal Transduction **(Root)**

R-HSA-9006931 Signaling by Nuclear Receptors **(Lineage)**

R-HSA-8939211 ESR-mediated signaling

R-HSA-9009391 Extra-nuclear estrogen signaling

R-HSA-9634638 Estrogen-dependent nuclear events downstream of ESR-membrane signaling **(Leaf)**

The second path comes from the Disease tree:

R-HSA-1643685 Disease **(Root)**

R-HSA-5663202 Diseases of signal transduction by growth factor receptors and second messengers **(Lineage)**

R-HSA-1227990 Signaling by ERBB2 in Cancer

R-HSA-9665686 Signaling by ERBB2 TMD/JMD mutants **(Leaf)**

As every path in a tree starts from the same root vertex, it is convenient to name each tree using the name of the root vertex. The procedure we have given decomposes *R*_*L*_ into 29 trees and a single crosstalk lattice summarizing all the crosstalk in *R*_*L*_. A list of all 29 trees named by root vertex (together with an entry for the crosstalk lattice) can be found in Table 1.

**Table 1.** Tree & Lattice Metrics.

| **Tree or Lattice** | $P$ | $\vert L\vert $ | $\vert \Pi\vert $ | $\vert V\vert $ | $D$ | $M$ | $C$ | $F$ | $\eta$ |
| --- | --- | --- | --- | --- | --- | --- | --- | --- | --- |
| Autophagy | 152 | 3 | 7 | 11 | 19 | 5 | 20 | 11 | U |
| Cell Cycle | 670 | 3 | 60 | 107 | 271 | 8 | 272 | 67 | 0.20 |
| Cell-Cell communication | 131 | 3 | 8 | 13 | 20 | 5 | 21 | 12 | 0.11 |
| Cellular responses to stimuli | 783 | 2 | 28 | 45 | 96 | 6 | 97 | 33 | 0.19 |
| Chromatin organization | 240 | 1 | 5 | 7 | 10 | 3 | 11 | 7 | U |
| Circadian Clock | 68 | 3 | 3 | 4 | 3 | 2 | 4 | 4 | U |
| DNA Repair | 315 | 7 | 40 | 64 | 131 | 7 | 132 | 46 | 0.21 |
| DNA Replication | 157 | 1 | 2 | 5 | 6 | 4 | 7 | 5 | U |
| Developmental Biology | 1134 | 12 | 51 | 68 | 160 | 6 | 161 | 56 | 0.11 |
| Digestion and absorption | 29 | 2 | 4 | 7 | 8 | 3 | 9 | 6 | 0.33 |
| Disease | 2169 | 12 | 540 | 701 | 2070 | 9 | 2071 | 548 | 0.10 |
| Drug ADME | 111 | 8 | 9 | 11 | 11 | 3 | 12 | 11 | U |
| Extracellular matrix organization | 300 | 9 | 12 | 19 | 22 | 4 | 23 | 15 | 0.50 |
| Gene expression (Transcription) | 1542 | 6 | 99 | 128 | 357 | 7 | 358 | 105 | 0.091 |
| Hemostasis | 726 | 7 | 23 | 34 | 53 | 4 | 54 | 26 | 0.29 |
| Immune System | 2265 | 3 | 122 | 172 | 427 | 7 | 428 | 128 | 0.15 |
| Metabolism | 2143 | 15 | 234 | 322 | 789 | 6 | 790 | 239 | 0.15 |
| Metabolism of RNA | 719 | 11 | 33 | 51 | 80 | 5 | 81 | 37 | 0.32 |
| Metabolism of proteins | 1950 | 8 | 79 | 112 | 269 | 6 | 270 | 84 | 0.15 |
| Muscle contraction | 204 | 3 | 9 | 11 | 16 | 3 | 17 | 11 | U |
| Neuronal System | 419 | 4 | 48 | 76 | 177 | 7 | 178 | 54 | 0.18 |
| Organelle biogenesis and maintenance | 298 | 2 | 9 | 13 | 22 | 4 | 23 | 12 | 0.091 |
| Programmed Cell Death | 216 | 2 | 26 | 42 | 98 | 7 | 99 | 32 | 0.15 |
| Protein localization | 166 | 4 | 4 | 6 | 5 | 3 | 6 | 6 | U |
| Reproduction | 114 | 1 | 3 | 5 | 6 | 3 | 7 | 5 | U |
| Sensory Perception | 642 | 4 | 10 | 17 | 23 | 4 | 24 | 13 | 0.36 |
| Signal Transduction | 2599 | 16 | 265 | 379 | 999 | 9 | 1000 | 273 | 0.15 |
| Transport of small molecules | 729 | 10 | 49 | 69 | 139 | 6 | 140 | 54 | 0.17 |
| Vesicle-mediated transport | 762 | 2 | 21 | 34 | 67 | 7 | 68 | 27 | 0.17 |
| Crosstalk Lattice* |  | 13 | 171 | 206 |  |  |  |  |  |

The metrics for the trees and crosstalk lattice. The metrics are defined in Sec. 4. The crosstalk lattice is included to provide an estimate of its size. It should be noted that the crosstalk lattice's lineages correspond to roots of trees. U indicates an undefined value as given in Eq. 10.

At this point we will assume that we can obtain a good approximationg of how a drug targets the Reactome while ignoring the crosstalk lattice and proceed to only use the trees in the sequel.

### 3.4 Augmented Trees & Augmented Reactome Data Function

Each vertex corresponds a pathways which consists of a set of interlinked reactions. We may then include in the data associated with each vertex the set of all proteins involved in the the corresponding set of reactions. By the construction, a child pathway contains a strict subset of the reactions of the parent pathway (i.e. the parent must have some reactions that are not in the child). This is just because there is no reason to give multiple names to the same pathway in the Reactome Knowledgebase. However, if we think of a vertex as the set of proteins involved in the reactions of the corresponding pathway, then it is possible for parent and child vertices to have the same set of proteins. We can express this for any vertex *V* having a non-empty set of child vertices 𝔙^′^(*V*) as:

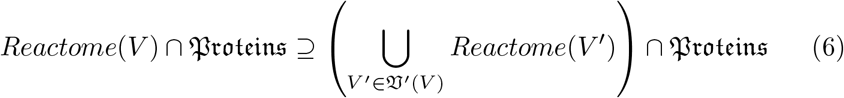

For the purpose of simplifying definitions and carrying out calculations on the trees, we would like Eq. 6 to be an equality. To make this an equality we require that every protein in a non-leaf vertex must appear one of its child vertices. This allows for us to perform the later bookkeeping of drug-target interactions in terms of paths as we have defined tham rather than forumulate a more complicated bookkeeping system where we have to account for reactions in non-leaf vertices that do not appear in leaves. We can accomplish this by constructing a set of 29 augmented trees from the 29 trees together with an augmented verson of *Reactome*(·).

Given a tree *T*, we construct the augmented tree *T*^*aug*^ and augmented function *Reactome*^*aug*^(*V*) by first making a copy of the tree. We then do the following:

For each vertex *V* ∈ *T* we:

1. Identify the set of all child vertices *V*^′^(*V*).
2. If 𝔙^′^(*V*)≠(∅ calculate the set) 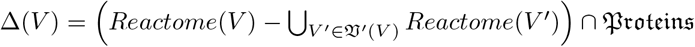
3. If Δ(*V*) is not empty we add an augment vertex *V* ^*aug*^ to the tree such that *V* ^*aug*^ is a child vertex of the vertex *V*. We further require: *Reactome*^*aug*^(*V* ^*aug*^) ∩ 𝔅 𝔯 𝔬 𝔱 𝔢 𝔦 𝔫 𝔰 = Δ(*V*)

The outcome of this procedure is illustrated for the toy model in Fig. 5. The corresponding paths are illustrated in Fig. 6. In constructing Fig.6 we assumed that every non-leaf vertex with only one child had proteins that were not present in the child vertices. This results in a doubling of the number of paths. The large ratio of regular paths to augment paths is due to the small number of regular paths. In practice, the ratio is much more favorable to regular paths as can be seen by comparing Tables 1 and 2

**Table 2.** Augmented Tree Metrics.

| Tree or Lattice | $P$ | $\vert L\vert $ | $\vert \Pi\vert $ | $\vert V\vert $ | $D$ | $M$ | $C$ | $F$ | $\eta$ |
| --- | --- | --- | --- | --- | --- | --- | --- | --- | --- |
| Autophagy | 152 | 3 | 8 | 12 | 21 | 5 | 22 | 12 | U |
| Cell Cycle | 670 | 4 | 77 | 124 | 343 | 8 | 344 | 84 | 0.15 |
| Cell-Cell communication | 131 | 3 | 10 | 15 | 27 | 5 | 28 | 14 | 0.071 |
| Cellular responses to stimuli | 783 | 2 | 37 | 54 | 132 | 6 | 133 | 42 | 0.13 |
| Chromatin organization | 240 | 1 | 6 | 8 | 12 | 3 | 13 | 8 | U |
| Circadian Clock | 68 | 4 | 4 | 5 | 4 | 2 | 5 | 5 | U |
| DNA Repair | 315 | 7 | 46 | 70 | 154 | 7 | 155 | 52 | 0.17 |
| DNA Replication | 157 | 2 | 5 | 8 | 12 | 4 | 13 | 8 | U |
| Developmental Biology | 1134 | 12 | 59 | 76 | 185 | 6 | 186 | 64 | 0.098 |
| Digestion and absorption | 29 | 2 | 5 | 8 | 10 | 3 | 11 | 7 | 0.25 |
| Disease | 2169 | 12 | 561 | 722 | 2163 | 9 | 2164 | 569 | 0.096 |
| Drug ADME | 111 | 8 | 9 | 11 | 11 | 3 | 12 | 11 | U |
| Extracellular matrix organization | 300 | 9 | 17 | 24 | 34 | 4 | 35 | 20 | 0.27 |
| Gene expression (Transcription) | 1542 | 6 | 105 | 134 | 383 | 7 | 384 | 111 | 0.084 |
| Hemostasis | 726 | 7 | 29 | 40 | 67 | 4 | 68 | 32 | 0.22 |
| Immune System | 2265 | 3 | 145 | 195 | 515 | 7 | 516 | 151 | 0.12 |
| Metabolism | 2143 | 15 | 275 | 363 | 944 | 6 | 945 | 280 | 0.12 |
| Metabolism of RNA | 719 | 11 | 36 | 54 | 88 | 5 | 89 | 40 | 0.29 |
| Metabolism of proteins | 1950 | 8 | 93 | 126 | 323 | 6 | 324 | 98 | 0.12 |
| Muscle contraction | 204 | 3 | 9 | 11 | 16 | 3 | 17 | 11 | U |
| Neuronal System | 419 | 4 | 60 | 88 | 229 | 7 | 230 | 66 | 0.13 |
| Organelle biogenesis and maintenance | 298 | 2 | 10 | 14 | 24 | 4 | 25 | 13 | 0.083 |
| Programmed Cell Death | 216 | 2 | 34 | 50 | 128 | 7 | 129 | 40 | 0.11 |
| Protein localization | 166 | 4 | 5 | 7 | 7 | 3 | 8 | 7 | U |
| Reproduction | 114 | 2 | 4 | 6 | 7 | 3 | 8 | 6 | U |
| Sensory Perception | 642 | 4 | 13 | 20 | 30 | 4 | 31 | 16 | 0.27 |
| Signal Transduction | 2599 | 17 | 320 | 434 | 1216 | 9 | 1217 | 328 | 0.12 |
| Transport of small molecules | 729 | 10 | 59 | 79 | 167 | 6 | 168 | 64 | 0.14 |
| Vesicle-mediated transport | 762 | 2 | 27 | 40 | 91 | 7 | 92 | 33 | 0.12 |

The metrics for the augmented trees. The metrics are defined in Sec. 4. U indicates an undefined value as given in Eq. 10.

**Figure 5.**
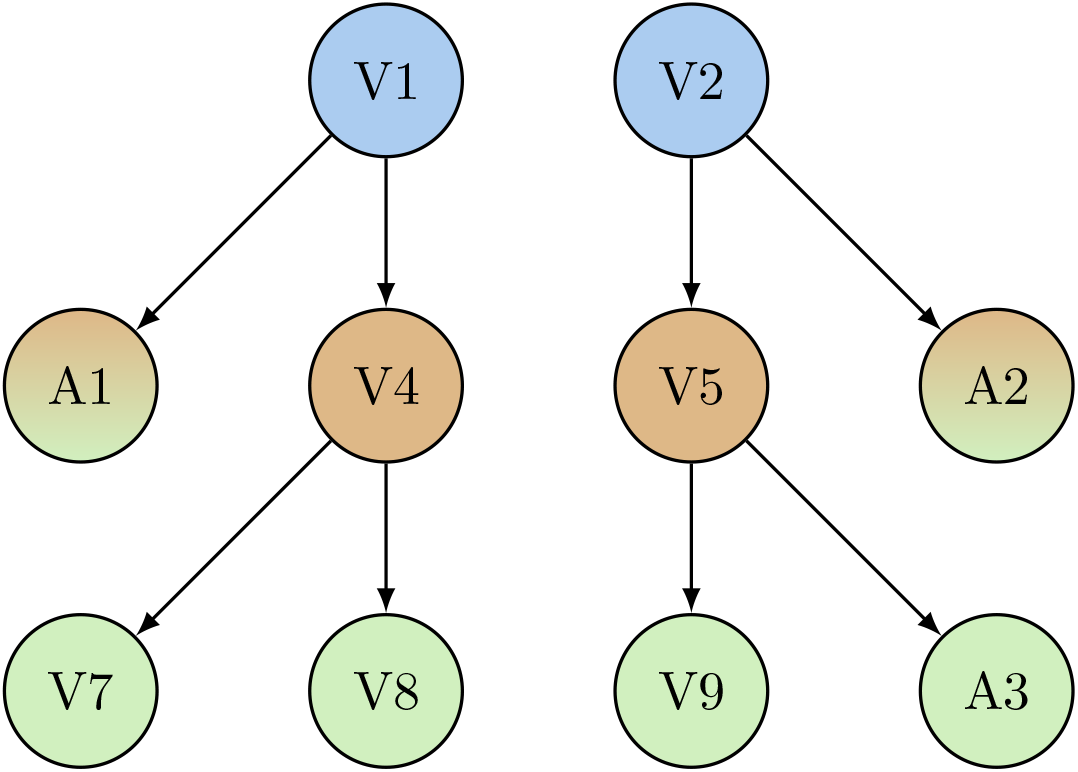
Toy Model Augmented Trees The augmented trees for the trees in Fig. 3. The augment vertices are labeled A1, A2, and A3. The blue vertices are the root vertices, the brown vertices are lineage vertices, the green vertices are the leaf vertices, the brown-green vertices are both lineage and leaf vertices. The colors of the non-augment vertices are the same as those in 3.

**Figure 6.**
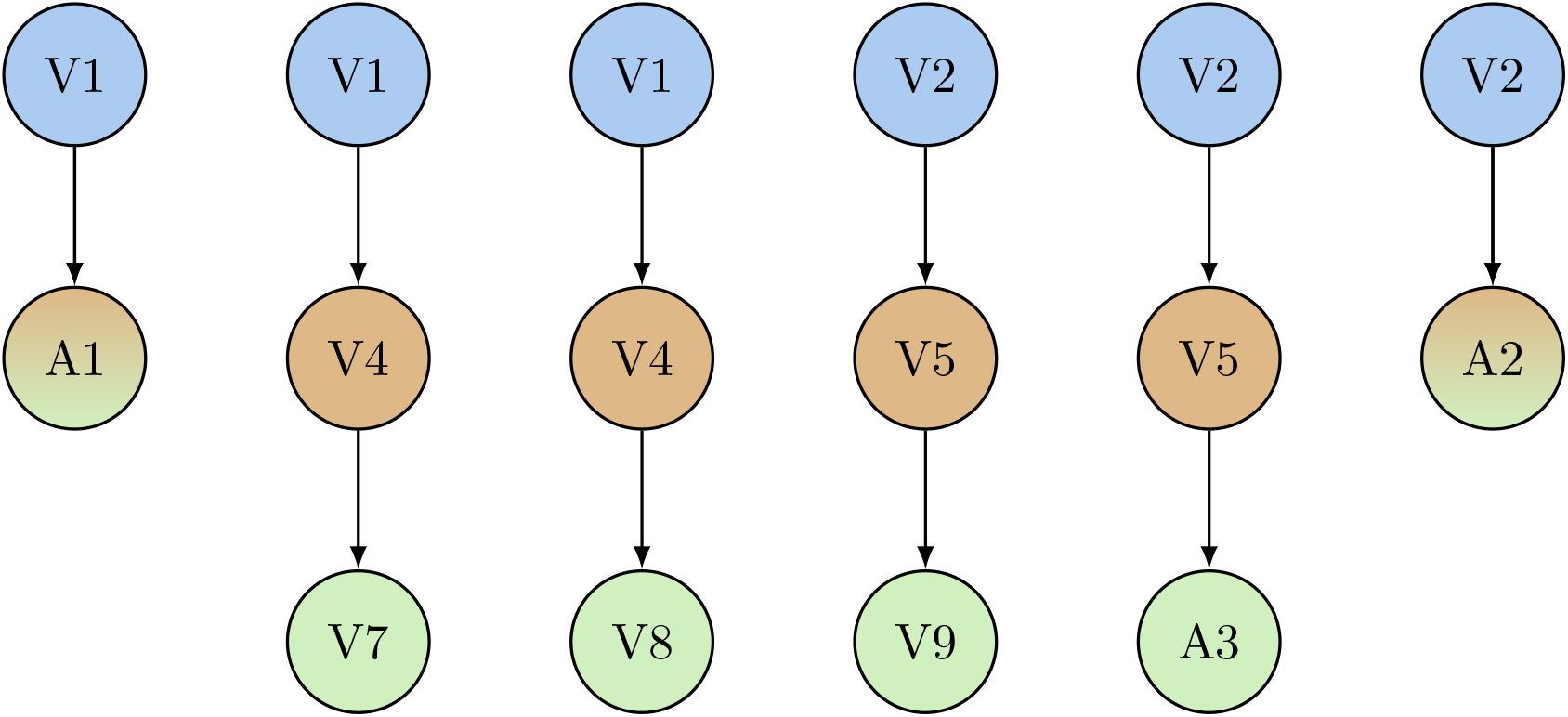
Toy Model Augmented Tree Paths The paths for the augmented trees in Fig. 5. The augment vertices are labeled A1, A2, and A3. The blue vertices are the root vertices, the brown vertices are lineage vertices, the green vertices are the leaf vertices, the brown-green vertices are both lineage and leaf vertices. The vertices are colored the same as in Fig. 5.

## 4 Metrics

At this point it is worthwhile to recall the original problem. The oncologist has identified a particular gene that they would like to target using a drug, and there are multiple candidate drugs that target the gene. The oncologist would like to choose from among the drugs the drug that has the greatest specificity and fewest off-target effects. In terms of the previous decompostion, this amounts to constructing for each drug the set of subtrees containing only those vertices containing proteins targeted by the drug, and comparing the sets to arrive at the best choice. Clearly this comparison is going to be a difficult task. We must therefore constuct metrics that measure the sizes and shapes of the subtrees in each set thereby reducing each set of subtrees to a set of metric values. Values for the metrics developed in this section are given for the regular tree and the crosstalk lattice in Table 1 and for the augmented tree in Table 2. The crosstalk lattice information has been retained in Table 1 to provide information as to how the crosstalk lattice compares in size to the trees.

### 4.1 Tree Structure Metrics

The first set of metrics that we construct aim to provide the clinician with a asummary of the geometric or topological shape or structure of a tree.

#### 4.1.1 Tree Size Metrics

We identify three metrics of the size of the tree. The first is the *number of lineages* |*L*| which is the number of child vertices of the root vertex of the tree. The second is the *number of paths* |Π| which is the total number of paths comprising the tree. The third is the *number of vertices* |*V*| which is the total number of vertices comprising the tree.

#### 4.1.2 Tree Conjointness Metric

The next property of a tree that is of interest for us is the conjointness of the tree. We motivate the idea of conjointness in the following way. Given a set of paths {Π} there are a number of trees that can be built using these paths. Each of these trees has its own number of vertices |*V*|. Clearly there is are minimum and maximum possible values of |*V*| corresponding to trees that can be constructed using the paths as well as a range of |*V*| for possible trees in between. We construct our conjointness metric with the aim of providing information as to where the actual tree being considered falls in this range given the set of paths {Π} that make up the tree.

We recall that the depth of a path is the total number of edges in the path. The *total path depth D* is the number we arrive at by summing the path depths of all the paths in the tree:

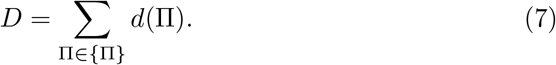

We begin by construcingt the *vertex ceiling C*. The vertex ceiling is the number of vertices in the largest tree that can contain all the paths in the original tree. The number of vertices in this tree is:

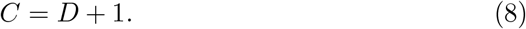

The *maximum path length M* which is the depth of the longest path in the tree.

Next we construct the *vertex floor F*. The vertex floor is then the number of vertices in the smallest tree that can contain all the paths in the original tree.

The number of vertices in this tree is:

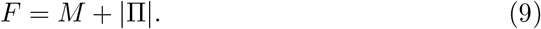

As expected *F* ≤ |*V*| ≤ *C*.

We can now construct the conjointness metric *η* as:

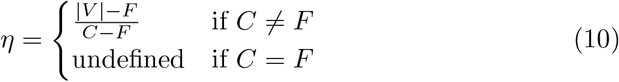

The ceiling, floor, and conjointness of a set of paths are illustrated in Fig. 7.

**Figure 7.**
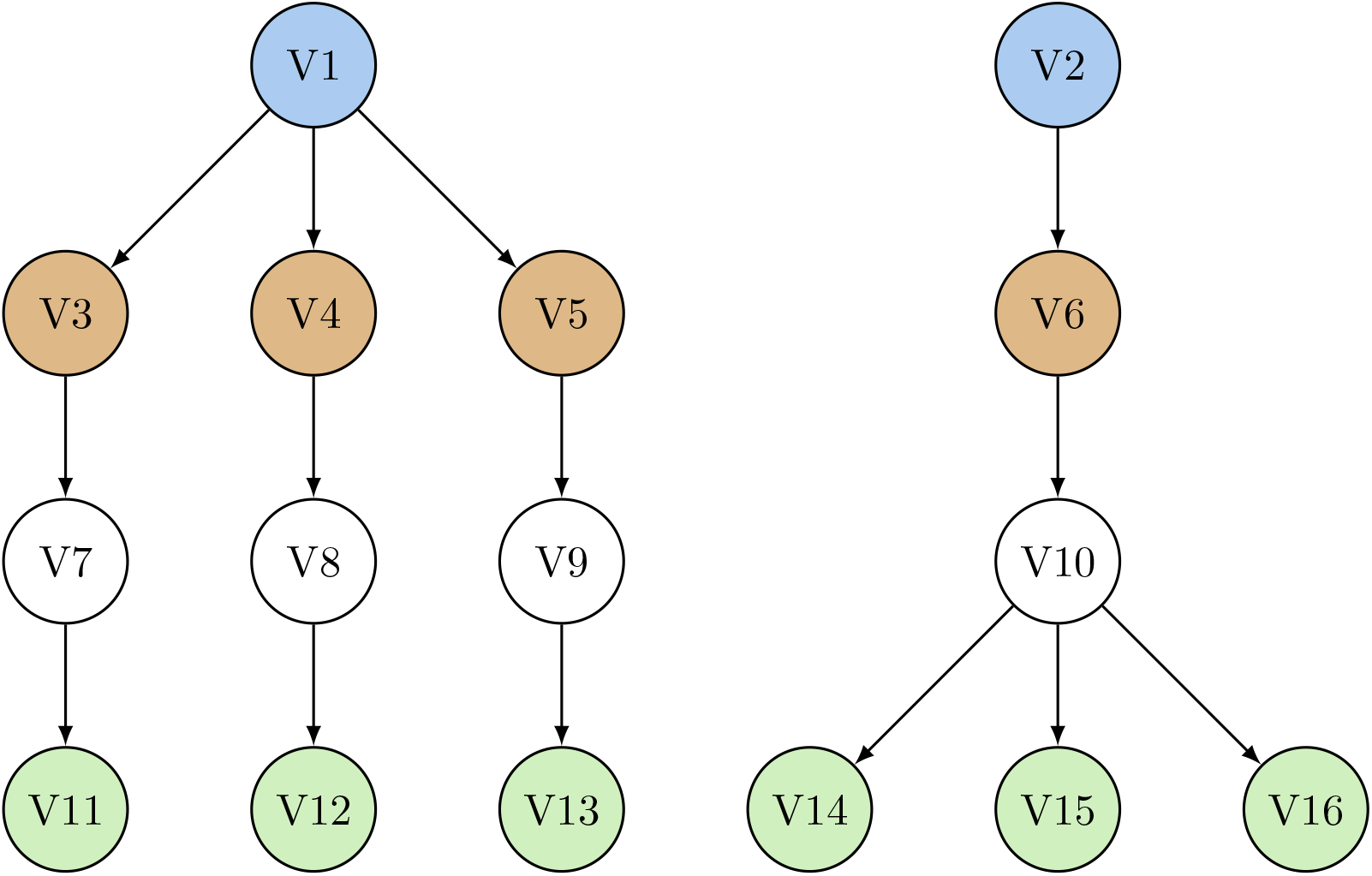
Conjointness Illustration for a Set of 3 Paths Each Having 4 Vertices Conjointness illustration for a set of 3 paths each having 4 vertices. The left tree gives the tree with the largest number of vertices for the given paths. The number of vertices it has is the vertex ceiling *C* = 10. Its conjointness is *η* = 1. The reight tree gives the tree with the smallest number of vertices for the given paths. The number of vertices it has is the vertex floor *F* = 6. Its conjointness is *η* = 0. Other tree arrangements of the 3 paths would have a number of vertices between 6 and 10, and 0 *< η <* 1. The blue vertices are the root vertices, the brown vertices are lineage vertices, the green vertices are the leaf vertex, the red vertex is a crosstalk vetex, the remaining vertices are white. The vertex labeling in this figure has no relationship to the vertex labeling in the previous figures.

### 4.2 Drug Coverage Metrics

We think of drug coverage in terms of how the pathways that are targeted by a drug relate to those pathways that are not targeted. We are given an augmented tree with size metrics *P*_*T*_, |*L*_*T*_ |, and |Π_*T*_ | and a conjointness metric *η*_*T*_, and a protein tree with size metrics |*L*_*P*_|, and |Π_*P*_| and a conjointness metric *η*_*P*_. We are also given the subtree consisting of vertices containing proteins that are targeted by the drug under consideration with size metrics *P*_*D*_, |*L*_*D*_|, and |Π_*D*_| and a conjointness metric *η*_*D*_e. We quantify this relationship in terms of a set of metrics which measure the fraction of an augmeted tree that is present in the subtree according to the metrics we ahve developed.

The first metric is the fraction of proteins in the augmented tree that are targeted by the drug. This is the *protein coverage metric K*_*P*_ :

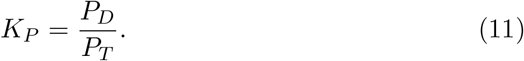

The second metric is the fraction of lineage pathways in the augmented tree that are targeted by the drug. This is the *tree lineage coverage metric K*_|_*L*|:

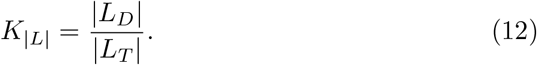

The third metric is the fraction of lineage pathways in the augmented tree that are targeted by the drug. This is the *protein lineage coverage metric k*_|_*L*|:

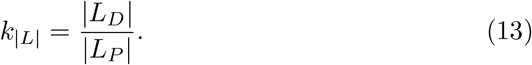

The fourth metric is the fraction of paths in the augmented tree that are targeted by the drug. This is the *tree path coverage metric K*_|_Π|:

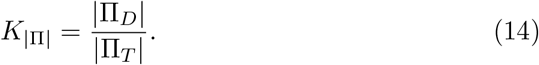

The fifth metric is the fraction of paths in the augmented tree that are targeted by the drug. This is the *protein path coverage metric k*_|_Π|:

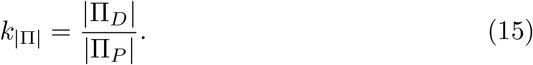

## 5 Empirical Studies

We illustrate the use of the metrics by examining the performance of the drugs Sunitinib Malate, Bosutinib, and Imatinib Mesylate in targeting EGFR (epiderimal growth factor receptor) which regulates cell growth, division, and survival. We consider 2 of the 29 trees dervied from the Reactome graph —the Signal Transduction tree and the Disease tree. These are the trees with the Signal Transduction pathway and Disease pathway as root pathways, respectively. The full list of trees named by root pathway can be found in Table 1 giving an idea what portion of the full Reactome is represented by the Signal Transduction and Disease trees. We are interested in how the metrics indicate the specificity and of-target effects of the 3 drugs.

For each drug and for each of the two trees we construct a subtree by identifying all the proteins targeted by the drug. In our construction we only include targets that have an evidence level of III in the Cancer Targetome, that is the drug-target relationship must have supporting literature information and at least one reported experimental binding value. Following [4], the maximum assay value was chosen to be 100 nM restricting the data to reactions that are relevant to drug binding [6, 7]. The data for this analysis are given in 4 tables. In Table 3 we give the metrics for subtrees constructed using only those pathways involving EGFR. In Tables 4, 5, and 6, we give the metrics for the subtrees constructed for Sunitinib Malate, Bosutinib, and Imatinib Mesylate, respectively.

**Table 3.** Protein Tree Metrics (EGFR)

| Tree or Lattice | $P$ | $\vert L\vert $ | $\vert \Pi\vert $ | $\vert V\vert $ | $D$ | $M$ | $C$ | $F$ | $\eta$ |
| --- | --- | --- | --- | --- | --- | --- | --- | --- | --- |
| Autophagy |  |  |  |  |  |  |  |  |  |
| Cell Cycle |  |  |  |  |  |  |  |  |  |
| Cell-Cell communication |  |  |  |  |  |  |  |  |  |
| Cellular responses to stimuli |  |  |  |  |  |  |  |  |  |
| Chromatin organization |  |  |  |  |  |  |  |  |  |
| Circadian Clock |  |  |  |  |  |  |  |  |  |
| DNA Repair |  |  |  |  |  |  |  |  |  |
| DNA Replication |  |  |  |  |  |  |  |  |  |
| Developmental Biology | 1134 | 1 | 1 | 5 | 4 | 5 | 5 | 5 | U |
| Digestion and absorption |  |  |  |  |  |  |  |  |  |
| Disease | 2169 | 2 | 8 | 18 | 27 | 5 | 28 | 12 | 0.38 |
| Drug ADME |  |  |  |  |  |  |  |  |  |
| Extracellular matrix organization |  |  |  |  |  |  |  |  |  |
| Gene expression (Transcription) | 1542 | 1 | 1 | 5 | 4 | 5 | 5 | 5 | U |
| Hemostasis |  |  |  |  |  |  |  |  |  |
| Immune System |  |  |  |  |  |  |  |  |  |
| Metabolism |  |  |  |  |  |  |  |  |  |
| Metabolism of RNA |  |  |  |  |  |  |  |  |  |
| Metabolism of proteins |  |  |  |  |  |  |  |  |  |
| Muscle contraction |  |  |  |  |  |  |  |  |  |
| Neuronal System |  |  |  |  |  |  |  |  |  |
| Organelle biogenesis and maintenance |  |  |  |  |  |  |  |  |  |
| Programmed Cell Death |  |  |  |  |  |  |  |  |  |
| Protein localization |  |  |  |  |  |  |  |  |  |
| Reproduction |  |  |  |  |  |  |  |  |  |
| Sensory Perception |  |  |  |  |  |  |  |  |  |
| Signal Transduction | 2599 | 7 | 18 | 42 | 61 | 6 | 62 | 23 | 0.49 |
| Transport of small molecules |  |  |  |  |  |  |  |  |  |
| Vesicle-mediated transport | 762 | 1 | 1 | 4 | 3 | 4 | 4 | 4 | U |

The metrics for EGFR. The metrics are defined in Sec. 4. U indicates an undefined value as given in Eq. 10.

**Table 4.** Drug Coverage Metrics (Sunitinib Malate)

| Tree | $P$ | $\vert L\vert $ | $\vert \Pi\vert $ | $\eta$ | $K_P$ | $K_{\vert L\vert }$ | $k_{\vert L\vert }$ | $K_{\vert \Pi\vert }$ | $k_{\vert \Pi\vert }$ | $\eta$ |
| --- | --- | --- | --- | --- | --- | --- | --- | --- | --- | --- |
| Autophagy | 152 | 3 | 8 | U | 0.026 | 0.33 |  | 0.38 |  | U |
| Cell Cycle | 670 | 4 | 77 | 0.15 | 0.010 | 0.50 |  | 0.21 |  | 0.40 |
| Cell-Cell communication | 131 | 3 | 10 | 0.071 | 0.0076 | 0.33 |  | 0.10 |  | U |
| Cellular responses to stimuli | 783 | 2 | 37 | 0.13 | 0.0089 | 0.50 |  | 0.11 |  | 0.43 |
| Chromatin organization | 240 | 1 | 6 | U |  |  |  |  |  |  |
| Circadian Clock | 68 | 4 | 4 | U | 0.029 | 0.25 |  | 0.25 |  | U |
| DNA Repair | 315 | 7 | 46 | 0.17 | 0.0095 | 0.14 |  | 0.087 |  | 0.12 |
| DNA Replication | 157 | 2 | 5 | U |  |  |  |  |  |  |
| Developmental Biology | 1134 | 12 | 59 | 0.098 | 0.0097 | 0.17 | 2.0 | 0.15 | 9.0 | 0.20 |
| Digestion and absorption | 29 | 2 | 5 | 0.25 |  |  |  |  |  |  |
| Disease | 2169 | 12 | 561 | 0.096 | 0.0065 | 0.25 | 1.5 | 0.12 | 8.6 | 0.18 |
| Drug ADME | 111 | 8 | 9 | U |  |  |  |  |  |  |
| Extracellular matrix organization | 300 | 9 | 17 | 0.27 | 0.0033 | 0.11 |  | 0.059 |  | U |
| Gene expression (Transcription) | 1542 | 6 | 105 | 0.084 | 0.012 | 0.17 | 1.0 | 0.17 | 18 | 0.15 |
| Hemostasis | 726 | 7 | 29 | 0.22 | 0.0028 | 0.29 |  | 0.069 |  | 1.0 |
| Immune System | 2265 | 3 | 145 | 0.12 | 0.0079 | 1.0 |  | 0.34 |  | 0.22 |
| Metabolism | 2143 | 15 | 275 | 0.12 | 0.0033 | 0.20 |  | 0.025 |  | 0.46 |
| Metabolism of RNA | 719 | 11 | 36 | 0.29 | 0.0056 | 0.091 |  | 0.028 |  | U |
| Metabolism of proteins | 1950 | 8 | 93 | 0.12 | 0.0026 | 0.25 |  | 0.054 |  | 0.50 |
| Muscle contraction | 204 | 3 | 9 | U | 0.020 | 0.67 |  | 0.44 |  | U |
| Neuronal System | 419 | 4 | 60 | 0.13 | 0.017 | 0.25 |  | 0.13 |  | 0.077 |
| Organelle biogenesis and maintenance | 298 | 2 | 10 | 0.083 | 0.010 | 1.0 |  | 0.20 |  | 1.0 |
| Programmed Cell Death | 216 | 2 | 34 | 0.11 | 0.0093 | 0.50 |  | 0.059 |  | 0.50 |
| Protein localization | 166 | 4 | 5 | U |  |  |  |  |  |  |
| Reproduction | 114 | 2 | 4 | U |  |  |  |  |  |  |
| Sensory Perception | 642 | 4 | 13 | 0.27 |  |  |  |  |  |  |
| Signal Transduction | 2599 | 17 | 320 | 0.12 | 0.017 | 0.76 | 1.9 | 0.23 | 4.2 | 0.30 |
| Transport of small molecules | 729 | 10 | 59 | 0.14 | 0.0014 | 0.10 |  | 0.017 |  | U |
| Vesicle-mediated transport | 762 | 2 | 27 | 0.12 | 0.0079 | 0.50 | 1.0 | 0.26 | 7.0 | 0.20 |

The metrics for Sunitinib Malate. The metrics are defined in Sec. 4. U indicates an undefined value as given in Eq. 10.

**Table 5.**
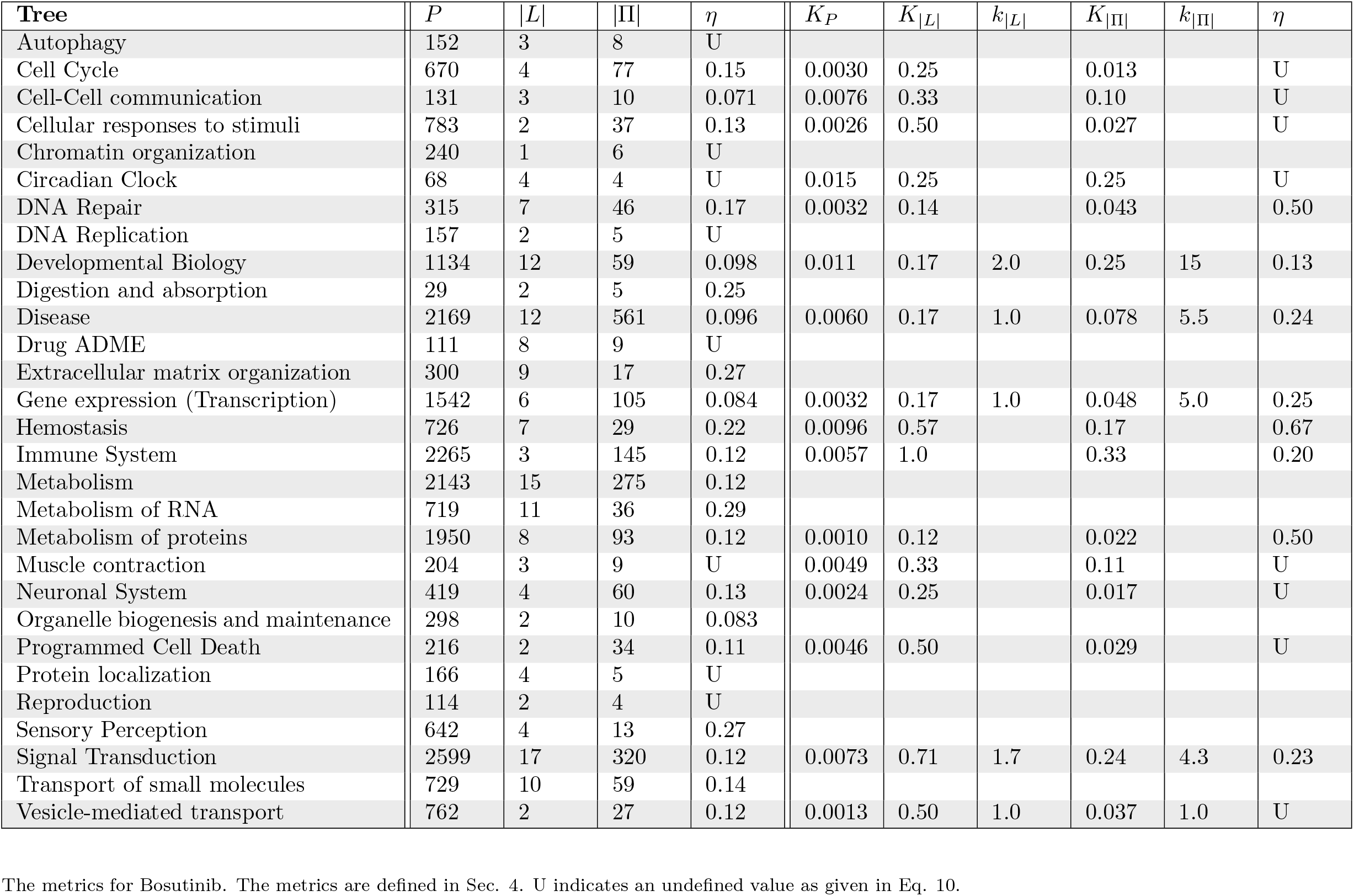
Drug Coverage Metrics (Bosutinib)

| Tree | $P$ | $\vert L\vert $ | $\vert \Pi\vert $ | $\eta$ | $K_P$ | $K_{\vert L\vert }$ | $k_{\vert L\vert }$ | $K_{\vert \Pi\vert }$ | $k_{\vert \Pi\vert }$ | $\eta$ |
| --- | --- | --- | --- | --- | --- | --- | --- | --- | --- | --- |
| Autophagy | 152 | 3 | 8 | U |  |  |  |  |  |  |
| Cell Cycle | 670 | 4 | 77 | 0.15 | 0.0030 | 0.25 |  | 0.013 |  | U |
| Cell-Cell communication | 131 | 3 | 10 | 0.071 | 0.0076 | 0.33 |  | 0.10 |  | U |
| Cellular responses to stimuli | 783 | 2 | 37 | 0.13 | 0.0026 | 0.50 |  | 0.027 |  | U |
| Chromatin organization | 240 | 1 | 6 | U |  |  |  |  |  |  |
| Circadian Clock | 68 | 4 | 4 | U | 0.015 | 0.25 |  | 0.25 |  | U |
| DNA Repair | 315 | 7 | 46 | 0.17 | 0.0032 | 0.14 |  | 0.043 |  | 0.50 |
| DNA Replication | 157 | 2 | 5 | U |  |  |  |  |  |  |
| Developmental Biology | 1134 | 12 | 59 | 0.098 | 0.011 | 0.17 | 2.0 | 0.25 | 15 | 0.13 |
| Digestion and absorption | 29 | 2 | 5 | 0.25 |  |  |  |  |  |  |
| Disease | 2169 | 12 | 561 | 0.096 | 0.0060 | 0.17 | 1.0 | 0.078 | 5.5 | 0.24 |
| Drug ADME | 111 | 8 | 9 | U |  |  |  |  |  |  |
| Extracellular matrix organization | 300 | 9 | 17 | 0.27 |  |  |  |  |  |  |
| Gene expression (Transcription) | 1542 | 6 | 105 | 0.084 | 0.0032 | 0.17 | 1.0 | 0.048 | 5.0 | 0.25 |
| Hemostasis | 726 | 7 | 29 | 0.22 | 0.0096 | 0.57 |  | 0.17 |  | 0.67 |
| Immune System | 2265 | 3 | 145 | 0.12 | 0.0057 | 1.0 |  | 0.33 |  | 0.20 |
| Metabolism | 2143 | 15 | 275 | 0.12 |  |  |  |  |  |  |
| Metabolism of RNA | 719 | 11 | 36 | 0.29 |  |  |  |  |  |  |
| Metabolism of proteins | 1950 | 8 | 93 | 0.12 | 0.0010 | 0.12 |  | 0.022 |  | 0.50 |
| Muscle contraction | 204 | 3 | 9 | U | 0.0049 | 0.33 |  | 0.11 |  | U |
| Neuronal System | 419 | 4 | 60 | 0.13 | 0.0024 | 0.25 |  | 0.017 |  | U |
| Organelle biogenesis and maintenance | 298 | 2 | 10 | 0.083 |  |  |  |  |  |  |
| Programmed Cell Death | 216 | 2 | 34 | 0.11 | 0.0046 | 0.50 |  | 0.029 |  | U |
| Protein localization | 166 | 4 | 5 | U |  |  |  |  |  |  |
| Reproduction | 114 | 2 | 4 | U |  |  |  |  |  |  |
| Sensory Perception | 642 | 4 | 13 | 0.27 |  |  |  |  |  |  |
| Signal Transduction | 2599 | 17 | 320 | 0.12 | 0.0073 | 0.71 | 1.7 | 0.24 | 4.3 | 0.23 |
| Transport of small molecules | 729 | 10 | 59 | 0.14 |  |  |  |  |  |  |
| Vesicle-mediated transport | 762 | 2 | 27 | 0.12 | 0.0013 | 0.50 | 1.0 | 0.037 | 1.0 | U |

The metrics for Bosutinib. The metrics are defined in Sec. 4. U indicates an undefined value as given in Eq. 10.

**Table 6.** Drug Coverage Metrics (Imatinib Mesylate)

| Tree | $P$ | $\vert L\vert $ | $\vert \Pi\vert $ | $\eta$ | $K_P$ | $K_{\vert L\vert }$ | $k_{\vert L\vert }$ | $K_{\vert \Pi\vert }$ | $k_{\vert \Pi\vert }$ | $\eta$ |
| --- | --- | --- | --- | --- | --- | --- | --- | --- | --- | --- |
| Autophagy | 152 | 3 | 8 | U |  |  |  |  |  |  |
| Cell Cycle | 670 | 4 | 77 | 0.15 | 0.0015 | 0.25 |  | 0.013 |  | U |
| Cell-Cell communication | 131 | 3 | 10 | 0.071 |  |  |  |  |  |  |
| Cellular responses to stimuli | 783 | 2 | 37 | 0.13 |  |  |  |  |  |  |
| Chromatin organization | 240 | 1 | 6 | U |  |  |  |  |  |  |
| Circadian Clock | 68 | 4 | 4 | U |  |  |  |  |  |  |
| DNA Repair | 315 | 7 | 46 | 0.17 | 0.0032 | 0.14 |  | 0.043 |  | 0.50 |
| DNA Replication | 157 | 2 | 5 | U |  |  |  |  |  |  |
| Developmental Biology | 1134 | 12 | 59 | 0.098 | 0.0026 | 0.17 | 2.0 | 0.051 | 3.0 | 0.33 |
| Digestion and absorption | 29 | 2 | 5 | 0.25 |  |  |  |  |  |  |
| Disease | 2169 | 12 | 561 | 0.096 | 0.0028 | 0.17 | 1.0 | 0.053 | 3.8 | 0.20 |
| Drug ADME | 111 | 8 | 9 | U |  |  |  |  |  |  |
| Extracellular matrix organization | 300 | 9 | 17 | 0.27 | 0.0067 | 0.11 |  | 0.059 |  | U |
| Gene expression (Transcription) | 1542 | 6 | 105 | 0.084 | 0.0026 | 0.17 | 1.0 | 0.038 | 4.0 | 0.25 |
| Hemostasis | 726 | 7 | 29 | 0.22 | 0.0028 | 0.43 |  | 0.10 |  | 1.0 |
| Immune System | 2265 | 3 | 145 | 0.12 | 0.0018 | 1.0 |  | 0.10 |  | 0.28 |
| Metabolism | 2143 | 15 | 275 | 0.12 |  |  |  |  |  |  |
| Metabolism of RNA | 719 | 11 | 36 | 0.29 |  |  |  |  |  |  |
| Metabolism of proteins | 1950 | 8 | 93 | 0.12 |  |  |  |  |  |  |
| Muscle contraction | 204 | 3 | 9 | U |  |  |  |  |  |  |
| Neuronal System | 419 | 4 | 60 | 0.13 |  |  |  |  |  |  |
| Organelle biogenesis and maintenance | 298 | 2 | 10 | 0.083 |  |  |  |  |  |  |
| Programmed Cell Death | 216 | 2 | 34 | 0.11 |  |  |  |  |  |  |
| Protein localization | 166 | 4 | 5 | U |  |  |  |  |  |  |
| Reproduction | 114 | 2 | 4 | U |  |  |  |  |  |  |
| Sensory Perception | 642 | 4 | 13 | 0.27 |  |  |  |  |  |  |
| Signal Transduction | 2599 | 17 | 320 | 0.12 | 0.0027 | 0.47 | 1.1 | 0.075 | 1.3 | 0.45 |
| Transport of small molecules | 729 | 10 | 59 | 0.14 | 0.0014 | 0.10 |  | 0.017 |  | U |
| Vesicle-mediated transport | 762 | 2 | 27 | 0.12 | 0.0013 | 0.50 | 1.0 | 0.037 | 1.0 | U |

The metrics for Imatinib Mesylate. The metrics are defined in Sec. 4. U indicates an undefined value as given in Eq. 10.

### 5.1 Signal Transduction Tree

The first tree dervided from the Reactome graph that we analyze is the Signal Transduction tree (i.e. the tree with the Signal Transduction pathway as the root pathway) which has as a natural function the prevention of cancer and is iinhibited in cancer. This is also the tree for which EGFR contains the most paths.

The simplest way to look at this would be to see if any of the proteins targeted by the either of the drugs appear in the biochemical reactions making up the tree. We assign the tree a value of 1 if it contains reactions involving any of the proteins targeted by the drug and a value of 0 otherwise. In this case, all three drugs target proteins in the Signal Transduction tree and thus we assign both drugs a value of 1. The most we can conclude from this information is that either drug is an equally good choice for treatment.

We might improve on this by looking at what fraction of the proteins present in the tree are targeted by the drugs. The Signal Transduction tree consists of *P* = 2599 proteins. We find that Sunitinib Malate targets *K*_*P*_ = 0.017 of the proteins, Bosutinib targets *K*_*P*_ = 0.0073, and Imatinib Mesylate targets *K*_*P*_ = 0.0027. Thus, ordering the drugs from greatest to least specificity we find Imatinib Mesylate, Bosutinib, Sunitinib Malate.

The first way we looked at the targeting of the tree by a drug was by simply asking whether the drug targeted any proteins in the tree. However this approach ignores available information about the reactions making up the tree and how they might be related to each other. We can try to obtain more information about how a drug targets the tree by grouping the reactions in the tree into disjoint sets and identifying which sets contain proteins targeted by the drug. The tree structure provides a guide for dividing the reactions into conceptually meaningful sets. We proceed by using the tree to construct the disjoint sets and then look at which sets are targeted by the drug.

The coarsest-grained division of reactions into sets that is available to us that obtained by looking at the lineage pathways of the tree. The Signal Transduction tree consists of |*L*| = 17 lineages, while the EGFR tree consists of |*L*| = 7 lineages. The Sunitinib Malate tree consists of |*L*| = 13 lineages, the Bosutinib tree of |*L*| = 12, and the Imatinib Mesylate tree of *L* = 8. Sunitinib Malate targets *K*_|*L*|_ = 13*/*17 = 0.76 of the lineages in the augmented tree, Bosutinib targets *K*_|*L*|_ = 12*/*17 = 0.71 of the lineages, and Imatinib Mesylate targets *K*_|*L*|_ = 8*/*17 = 0.47 of the lineages. Sunitinib Malate targets *k*_|*L*|_ = 13*/*7 = 1.9 of the lineages in the protein tree, Bosutinib targets *k*_|*L*|_ = 12*/*7 = 1.7 of the lineages, and Imatinib Mesylate targets *k*_|*L*|_ = 8*/*7 = 1.1 of the lineages. Thus, ordering the drugs from greatest to least specificity we find Imatinib Mesylate, Bosutinib, Sunitinib Malate.

The finest-grained division of reactions into sets is at the level of leaf pathways. In the analysis of the tree that we have carried out, every leaf is associated with a unique path, so we can proceed by looking at paths. The Signal Transduction tree consists of |Π| = 320 paths, while the Sunitinib Malate tree consists of |Π| = 75, the Bosutinib tree of |Π| = 78 paths, and the Imatinib Mesylate tree of |Π| = 24 paths. Sunitinib Malate targets *K*_|Π|_ = 75*/*320 = 0.23 of the paths in the augmented tree, Bosutinib targets *K*_|Π|_ = 78*/*320 = 0.24 of the paths, and Imatinib Mesylate targets *K*_|Π|_ = 24*/*320 = 0.075 of the paths. Sunitinib Malate targets *k*_|Π|_ = 75*/*18 = 4.2 of the paths in the protein tree, Bosutinib targets *k*_|Π|_ = 78*/*18 = 4.3 of the paths, and Imatinib Mesylate targets *k*_|Π|_ = 24*/*18 = 1.3 of the paths. Thus, ordering the drugs from greatest to least specificity we find Imatinib Mesylate, Sunitinib Malate, Bosutinib.

The conjointness metric *η* as provides information about how quickly the tree branches out from the lineage vertices to the leaf vertices. Conjointness values closer to 1 indicate that the bulk of the branching out happens closer to the lineage vertices while values closer to 0 indicate that the bulk of the branching outn happens closer to the leaf vertices. The Signal Transduction tree branches out from |*L*| = 17 to |Π| = 320 with a conjointness of *η* = 0.12, while the Sunitinib Malate tree branches out from |*L*| = 13 to |Π| = 75 with a conjointness of *η* = 0.30, Bosutinib tree branches out from |*L*| = 12 to |Π| = 78 with a conjointness of *η* = 0.23, and the Imatinib Mesylate tree branches out from |*L*| = 8 to |Π| = 24 with a conjointness of *η* = 0.45. We see that Signal Transduction tree does most of its branching out nearer to the leaves, while the drugs do most of their branching earlier on.

### 5.2 Disease Tree

The second tree dervided from the Reactome graph that we analyze is the Disease tree (i.e. the tree with the Disease pathway as the root pathway). We proceed with the same analysis as we carried out for the Signal Transduction tree.

We again begin by looking to see if any of the proteins targeted by the either of the drugs appear in the biochemical reactions making up the tree. In this case, all three drugs target proteins in the Disease tree and thus we assign both drugs a value of 1. The most we can conclude from this information is that either drug is an equally good choice for treatment.

We again move by looking at what fraction of the proteins present in the tree are targeted by the drugs. The Disease tree consists of *P* = 2599 proteins. We find that Sunitinib Malate targets *K*_*P*_ = 0.0065 of the proteins, Bosutinib targets *K*_*P*_ = 0.0060, and Imatinib Mesylate targets *K*_*P*_ = 0.0028. Thus, ordering the drugs from greatest to least specificity we find Imatinib Mesylate, Bosutinib, Sunitinib Malate.

jWe again proceed to lookin at the lineage pathways of the tree. The Disease tree consists of *L* = 12 lineages, while the EGFR tree consists of |*L*| = 2 lineages. The Sunitinib Malate tree consists of *L* = 3 lineages, the Bosutinib tree of *L* = 2, and the Imatinib Mesylate tree of *L* = 2. Sunitinib Malate targets *K*_|*L*|_ = 3*/*12 = 0.25 of the lineages in the augmented tree, Bosutinib targets *K*_|*L*|_ = 2*/*12 = 0.17 of the lineages, and Imatinib Mesylate targets *K*_|*L*|_ = 2*/*12 = 0.17 of the lineages. Thus drugs Bosutinib and Imatinib Mesylate have the same and lowest specificity while Sunitinib Malate has less specificity. Sunitinib Malate targets *k*_|*L*|_ = 3*/*2 = 1.5 of the lineages in the protein tree, Bosutinib targets *k*_|*L*|_ = 2*/*2 = 1.0 of the lineages, and Imatinib Mesylate targets *k*_|*L*|_ = 2*/*2 = 1.0 of the lineages. Thus drugs Bosutinib and Imatinib Mesylate have the same and lowest specificity while Sunitinib Malate has less specificity.

We conntinue by looking again at the level of leaf pathways. The Disease tree consists of |Π| = 561 paths, while the Sunitinib Malate tree consists of |Π| = 69, the Bosutinib tree of |Π| = 44 paths, and the Imatinib Mesylate tree of |Π| = 30 paths. Sunitinib Malate targets *K*_|Π|_ = 69*/*561 = 0.12 of the paths in the augmented tree, Bosutinib targets *K*_|Π|_ = 44*/*561 = 0.078 of the paths, and Imatinib Mesylate targets *K*_|Π|_ = 30*/*561 = 0.053 of the paths. Sunitinib Malate targets *k*_|Π|_ = 69*/*8 = 8.6 of the paths in the protein tree, Bosutinib targets *k*_|Π|_ = 44*/*8 = 5.5 of the paths, and Imatinib Mesylate targets *k*_|Π|_ = 30*/*8 = 3.8 of the paths.Thus, ordering the drugs from greatest to least specificity we find Imatinib Mesylate, Bosutinib, Sunitinib Malate.

Finally, we look again at the conjointness. The Disease tree branches out from |*L*| = 12 to |Π| = 561 with a conjointness of *η* = 0.096, while the Sunitinib Malate tree branches out from |*L*| = 3 to |Π| = 69 with a conjointness of *η* = 0.18, Bosutinib tree branches out from |*L*| = 2 to |Π| = 44 with a conjointness of *η* = 0.24, and the Imatinib Mesylate tree branches out from |*L*| = 2 to |Π| = 30 with a conjointness of *η* = 0.20. We see that Disease tree does most of its branching out nearer to the leaves, while the drugs do most of their branching earlier on.

## 6 Discussion

The original problem was to characterize a drug ”promiscuity”, that is its targeting of multiple genes or pathways beyond those to be targeted in a specific treatment. We would like a drug to exhibit high specificity (targeting few genes or pathways beyond those that are immediately relevant) and few off-target effects. We have captured measures which provide information about these characteristics in terms of a set 29 trees that we extracted out of the Reactome Knowledgebase for each of which we construct 2 metrics, |*L*| and |Π|, that characterize the size of the tree, a metric, *η*, that characterizes the shape of a tree, and 4 metrics, *K*_|*L*|_, *k*_|*L*|_, *K*_|Π|_, and *k*_|Π|_, which express the fraction of a tree that is targeted by the drug. We use the two metrics, |*L*| and |Π|, to provide an indication of the size of the original tree and how it compares to the other 28 trees in size, and to use the 4 metrics, *K*_|*L*|_, *k*_|*L*|_, *K*_|Π|_, and *k*_|Π|_, to indicate what fraction of the tree is being tageted by the drug. The remaining metric, *η*, provides the oncologist with more fine-grained information about the tree structure that may prove useful. In doing so, we have reduced the large and complex Reactome graph and information about drug-target interactions to a more manageable set of metric values that admit straightforward numerical comparisons. The number of metrics is nevertheless still quite large and likely the oncologist will find, using our metrics, that a situation may arise where one candidate drug exhibits higher specificity on some set of trees, while another candidate exhibits higher specificity on another. This sort of problem must be resolved through some scheme of weighing the relative importance of the trees that is beyond the scope of the present work.

## References

1. Siegel, R. L., Miller, K. D., Fuchs, H. E. & Jemal, A. Cancer statistics, 2022. CA: A Cancer Journal for Clinicians 72, 7–33. eprint: https://acsjournals.onlinelibrary.wiley.com/doi/pdf/10.3322/caac.21708. https://acsjournals.onlinelibrary.wiley.com/doi/abs/10.3322/caac.21708 (2022).

2. Akhoon, N. Precision Medicine: A New Paradigm in Therapeutics. International journal of preventive medicine 12. eprint: 10.4103%2Fijpvm.IJPVM_375_19. 10.4103%2Fijpvm.IJPVM_375_19 (Feb. 2021).

3. Gillespie, M. et al. The reactome pathway knowledgebase 2022. Nucleic Acids Research 50, D687–D692. issn: 0305-1048. eprint: https://academic.oup.com/nar/article-pdf/50/D1/D687/42058295/gkab1028.pdf. 10.1093/nar/gkab1028 (Nov. 2021).

4. Blucher, A. S., Choonoo, G., Kulesz-Martin, M., Wu, G. & McWeeney, S. K. Evidence-Based Precision Oncology with the Cancer Targetome. Trends in Pharmacological Sciences 38, 1085–1099. issn: 0165-6147. https://www.sciencedirect.com/science/article/pii/S0165614717301669 (2017).

5. Reactome Data Model Accessed on November 13, 2023. https://reactome.org/documentation/data-model.

6. Paolini, G. V., Shapland, R. H. B., van Hoorn, W. P., Mason, J. S. & Hopkins, A. L. Global mapping of pharmacological space. Nature biotechnology 24, 805–815 (July 2006).

7. Wang, Y. et al. Evidence-Based and Quantitative Prioritization of Tool Compounds in Phenotypic Drug Discovery. Cell chemical biology 23, 862– 874 (July 2016).

